# A conditioned place preference for heroin is signaled by increased dopamine and direct pathway activity and decreased indirect pathway activity in the nucleus accumbens

**DOI:** 10.1101/2021.07.14.452340

**Authors:** Timothy J. O’Neal, Mollie X. Bernstein, Derek J. MacDougall, Susan M. Ferguson

## Abstract

Initial drug use promotes the development of conditioned reinforcement, whereby the reinforcing properties of a drug become attributed to drug-associated stimuli, such as cues and contexts. A principal role for the nucleus accumbens (NAc) in the response to drug-associated stimuli has been well-documented. In particular, direct and indirect pathway medium spiny neurons (dMSNs and iMSNs) have been shown to bidirectionally regulate cue-induced heroin-seeking in rats expressing addiction-like phenotypes, and a shift in NAc activity towards the direct pathway has been shown in mice following cocaine conditioned place preference (CPP). However, how NAc signaling guides heroin CPP, and whether heroin alters the balance of signaling between dMSNs and iMSNs remains unknown. Moreover, the role of NAc dopamine signaling in heroin reinforcement remains unclear. Here, we integrate fiber photometry for *in vivo* monitoring of dopamine and dMSN/iMSN calcium activity with a heroin CPP procedure in rats to address these outstanding questions. We identify a sensitization-like response to heroin in the NAc, with prominent iMSN activity during initial heroin exposure and prominent dMSN activity following repeated heroin exposure. We demonstrate a ramp in dopamine activity, dMSN activation, and iMSN inactivation preceding entry into a heroin-paired context, and a decrease in dopamine activity, dMSN inactivation, and iMSN activation preceding exit from a heroin-paired context. Finally, we show that buprenorphine is sufficient to prevent the development of heroin CPP and activation of the NAc post-conditioning. Together, these data support the hypothesis that an imbalance in NAc activity contributes to the development of addiction.

**SIGNIFICANCE STATEMENT:** The attribution of the reinforcing effects of drugs to neutral stimuli (e.g., cues and contexts) contributes to the maintenance of addiction, as re-exposure to drug-associated stimuli can reinstate drug seeking and taking even after long periods of abstinence. The nucleus accumbens (NAc) has an established role in encoding the value of drug-associated stimuli, and dopamine release into the NAc is known to modulate the reinforcing effects of drugs, including heroin. Using fiber photometry, we show that entering a heroin-paired context is driven by dopamine signaling and NAc direct pathway activation, whereas exiting a heroin-paired context is driven by NAc indirect pathway activation. This study provides further insight into the role of NAc microcircuitry in encoding the reinforcing properties of heroin.

## INTRODUCTION

The opioid epidemic remains a major public health crisis in the United States, with relapse rates and overdose-related fatalities continuing to rise (Hedegaard et al., 2018). One critical mechanism underlying persistent drug-craving is conditioned drug reinforcement: the attribution of the reinforcing properties of a drug to a previously neutral stimulus (conditioned stimulus (CS); e.g., cues, contexts) (Everitt et al., 2018). Re-exposure to a drug-paired CS can reinstate drug-seeking following action/outcome devaluation (Shaham et al., 2003) or prolonged abstinence (Grimm et al., 2001). The nucleus accumbens (NAc) integrates cortical and subcortical inputs to guide behavioral processes associated with addiction, including associative learning, decision making, and motivation (Gerfen and Surmeier, 2011; Calabresi et al., 2014; Koob and Volkow, 2016). The NAc is heterogeneous, and predominantly comprised of two interspersed populations of medium spiny neurons (MSNs): direct pathway MSNs (dMSNs) that project to the ventral tegmental area (VTA) and primarily express the dopamine (DA) D1 receptor, and indirect pathway MSNs (iMSNs) that project to the ventral pallidum (VP) and primarily express the DA D2 receptor. These cell populations can have opposing control over behavioral output, with dMSNs serving as a “go” signal to facilitate behavioral actions and iMSNs serving as a “stop” signal to suppress or terminate unwanted actions (Albin et al., 1989; Kravitz et al., 2010; Cui et al., 2013; Macpherson et al., 2014). Moreover, emerging evidence has demonstrated a role for NAc dMSNs and iMSNs in encoding variability to the addictive effects of illicit drugs (Volkow et al., 2002; Nader et al., 2006; Belin et al., 2008; Bock et al., 2013; Yager et al., 2019; O’Neal et al., 2020), highlighting a central role for NAc MSNs in the transition to addiction.

Midbrain DA release into the NAc is central to the rewarding effects of illicit drugs (Crummy et al., 2020), and phasic DA release into the NAc promotes drug-seeking (Phillips et al., 2003). In addition to slower modulation of dMSNs and iMSNs via activation of G-protein coupled D1 and D2 receptors, DA can rapidly modulate these neuronal populations via opening of IP_3_ receptors and release of intracellular Ca^2+^ (Swapna et al., 2016), a process that is integral for the propagation of Ca^2+^ waves and heterosynaptic plasticity in the NAc (Bailey et al., 2000; Plotkin et al., 2013). *In vivo* imaging of dMSN and iMSN Ca^2+^ activity has revealed a rapid increase in dMSN activity and a progressive decrease in iMSN activity after acute cocaine exposure (Luo et al., 2011), as well as a persistent attenuation of iMSN activity after chronic cocaine treatment (Park et al., 2013); all of which lead to a long-lasting predominance of dMSN over iMSN signaling. Moreover, a ramping of dMSN Ca^2+^ activity along with a concomitant decrease in iMSN activity has been observed in mice preceding entry into a context associated with cocaine treatment (Calipari et al., 2016). Importantly, psychostimulants and opioids have different mechanisms of action and engage non-overlapping subcircuits to promote addictive behaviors (Badiani et al., 2011; Crummy et al., 2020), so the temporal relationship between NAc signaling and heroin-seeking remains unknown.

While targeted manipulations of dMSNs and iMSNs have demonstrated oppositional control over addictive behaviors (Lobo et al., 2010; Ferguson et al., 2011; O’Neal et al., 2020), it remains to be determined how the acquisition of a heroin CS is encoded by DA, dMSN, and iMSN signaling in the NAc. Thus, we combined fiber photometry for *in vivo* monitoring of NAc DA signaling and dMSN/iMSN Ca^2+^ signaling with a heroin conditioned place preference (CPP) procedure. NAc activity was recorded during early and late conditioning sessions to assess changes in the neural response over the course of conditioning. Following conditioning, we examined temporally precise DA signaling and activity of dMSNs and iMSNs during entries to and exits from contexts that had been paired with either saline or heroin and compared these signals with those recorded pre-conditioning. Finally, we explored the effects of buprenorphine, a partial mu opioid receptor agonist used in opioid-replacement therapy, on the acquisition of heroin CPP and subsequent activation of the NAc.

## MATERIALS AND METHODS

### Subjects

Outbred female (*n* = 40; ~8 weeks old, 175-199 g at arrival; Envigo) and male (*n* = 40; ~8 weeks old, 250-274 g at arrival; Envigo) Sprague-Dawley rats were pair-housed in a humidity- and temperature-controlled vivarium, with *ad libitum* access to food and water throughout the experiments. Rats were acclimated to the vivarium for at least five days and handled for at least three days prior to any procedures. All procedures were done in accordance with the National Institutes of Health’s Office of Laboratory Animal Welfare and were approved by the Seattle Children’s Research Institute’s Institutional Animal Care and Use Committee.

### Drugs

Diamorphine HCl (heroin) was obtained through the Drug Supply Program of the National Institute on Drug Abuse (NIDA) and was dissolved in sterile saline (0.9%) to a concentration of 1-3 mg/ml and administered at a dose of 1 ml/kg. Buprenorphine was also obtained through the Drug Supply Program of NIDA and was dissolved in sterile saline (0.9%) to a concentration of 0.2 mg/ml and administered at a dose of 1 ml/kg.

### Viral vectors

Adeno-associated viruses containing Flp recombinase (AAVrg-EF1a-flpo; #55637-AAVrg), Flp-dependent GCaMP6s (AAV8-EF1a-fDIO-GCaMP6s; #105714-AAV8), and Cre-dependent RCaMP1b (AAV1-Syn-FLEX-NES-jRCaMP1b-WPRE-SV40; #100850-AAV1) were acquired from Addgene and had titers of ≥1×10^13^ viral genomes/ml. dLight1.3b plasmid (AAV1-Syn-dLight1.3b) was acquired from Addgene (#135762) and prepared by the UW Molecular Genetics Resource Core. Canine adenovirus containing Cre recombinase (CAV2-Cre) had a titer of ~2.5×10^9^ viral genomes/*μ*l and was prepared as previously described (Kremer et al., 2009).

### Stereotaxic surgery

Rats were anesthetized with isoflurane (3% induction, 1-2% maintenance; Patterson Veterinary) and injected with meloxicam (1 mg/ml, 1 ml/kg *sc*; Patterson Veterinary) for analgesia. Following head-fixation in a digital stereotax (Kopf Instruments), the skull was exposed and scored, and craniotomies were drilled above target nuclei. Viral vectors were loaded into 10 *μ*l gas-tight syringes (Hamilton Company) and infused unilaterally into target nuclei (500 nl per virus, 100 nl/min). Coordinates (in mm, relative to Bregma (Paxinos et al., 1980)) were as follows: NAc [A/P +1.5, M/L +1.0, D/V −7.5], VP [A/P +0.2, M/L +2.0, D/V − 8.0], VTA [A/P −5.3, M/L +0.8 D/V −8.3]. To target dMSNs, AAVrg-EF1a-flpo was infused into the VTA and AAV8-EF1a-fDIO-GCaMP6s was infused into the NAc. To target iMSNs, CAV2-Cre was infused into the VP and AAV1-Syn-FLEX-NES-jRCaMP1b-WPRE-SV40 was infused into the NAc. Syringes were left in place for an additional 5 min following infusion and slowly retracted to allow proper diffusion of virus into target nuclei. Following viral infusion, fiber optic cannula (MFC_400/430-0.37_8mm_ZF2.5_FLT; >85% transmittance; Doric Lenses) were implanted in the NAc medial shell [D/V −7.4] and secured with skull screws, metabond (Patterson Dental), and dental cement.

### Fiber photometry

Three weeks after surgery, fiber photometry recordings were performed using a multiplex photometry system (FP3001; Neurophotometrics Ltd., San Diego, CA). Rats were acclimated to branching fiber optic patch cords (BFP4_400/440/LWMJ-0.37_3m_SMA*-4xFC; Doric Lenses) for at least three days prior to testing, and biosensor expression was confirmed the day prior to testing via brief (~5 min) home cage recordings. For all recordings, LEDs were heated at 100% power for at least 5 min prior to recording to minimize heat-induced LED decay during recordings, then reduced to <50 *μ*W for the duration of recordings (415 nm and 470 nm: ~12.5 *μ*W; 560 nm: ~25 *μ*W). Photometry recordings with a single biosensor (dLight1.3b imaging) used 2-phase cycling of 415 nm and 470 nm LEDs, and recordings with two biosensors (dual GCaMP6s/RCaMP1b imaging) used 3-phase cycling of 415 nm, 470 nm, and 560 nm LEDs. Fluorescent signals were bandpass filtered, collimated, reflected by a dichroic mirror, and focused by a 20x objective (0.40 NA) on the sensor of a CMOS camera. Signals were imaged at ~40 Hz, collected via custom-written workflows in Bonsai, and exported for off-line analysis via custom-written Python scripts.

### Conditioned place preference

#### CPP apparatus

Behavioral sessions were conducted in custom-built acrylic chambers (TAP Plastics, Seattle, WA) with two equally sized but visually distinct chambers (15” wide × 15” long × 12” tall), and a smaller center chamber (6” wide × 15” long × 12” tall) separated from the outer chambers via removable guillotine doors (3” wide × 12” tall). The walls of the two outer chambers were covered with visually distinct wallpapers (honeycomb and zebra, both grayscale), and the floor of the entire apparatus was matte black to facilitate behavioral tracking. USB cameras (2.1mm, wide-angle; ELP) were positioned above CPP chambers and interfaced with Bonsai for motion tracking. Video streams were greyscaled and thresholded, and binary region analysis was used to detect animal position within the CPP chambers. Chambers were cleaned between sessions with 70% ethanol (w/v).

#### CPP procedure

Experimental parameters were chosen based on a meta-analysis of heroin CPP by Bardo et al., 1995 to maximize potential effect sizes. The overall CPP procedure included an initial preference test (15 min, beginning at 11:00), eight conditioning sessions (2x/day, 40 min each, beginning at 09:00 and 13:00), and a final preference test (15 min, beginning at 11:00). During the preference tests, the guillotine doors were removed, and subjects were placed in the center of the neutral chamber and allowed to freely explore the full apparatus. During conditioning sessions, subjects received *ip* injections of saline (morning) or heroin (afternoon) and were immediately confined to one of the outer chambers. Saline and heroin pairings were assigned in a pseudorandomized, counterbalanced design to avoid sex or treatment biases with either chamber. Pretreatment injections of buprenorphine were given 10 min prior to injections of saline or heroin, when appropriate. All behavioral testing was done during the light cycle, due to the use of visual cues for conditioning.

### Immunohistochemistry

After behavioral testing, rats were deeply anesthetized with Euthasol (2 ml/kg *ip*; 3.9 mg/ml pentobarbital sodium and 0.5 mg/ml phenytoin sodium; Patterson Veterinary) and transcardially perfused with phosphate-buffered saline (PBS; pH = 7.4) followed by paraformaldehyde (PFA; 4% in PBS). Brains were extracted, fixed overnight in 4% PFA, post-fixed for >48 h in sucrose (30% in PBS), and sectioned (50 *μ*m) with a vibrating microtome. Floating sections were washed (PBS; 3×10 min), blocked (0.25% Triton-X, 5% normal goat serum, PBS; 2h), and incubated with primary antibodies (0.25% Triton-X, 2.5% normal goat serum, PBS; 24h) against Fos (1:800 rabbit anti-cFos, Cell Signaling #2250; RRID: AB_2247211) or tyrosine hydroxylase (1:400 mouse anti-TH, Sigma #MAB318; RRID: AB_2201528). Sections were then washed (PBS; 3×10 min) and incubated with secondary antibodies (0.25% Triton-X, 2.5% normal goat serum, PBS; 2h) conjugated to AF-568 (1:500 goat-anti-rabbit, Life Technologies #A11011; RRID: AB_143157) or AF-647 (1:500 goat-anti-mouse, Life Technologies #A21236; RRID: AB_2535805). Finally, sections were washed (PBS; 3×10min), mounted on slides, and cover-slipped with mounting medium with DAPI (Vectashield). Z-stacks along the rostral/caudal axis of the NAc (A/P +2.5 through A/P +0.7) were collected with confocal microscopy (20x; Zeiss LSM 710), and Fos^+^ cells were quantified using ImageJ (V1.49; NIH) and Adobe Illustrator (CC 2020).

### Experimental design and statistical analyses

Behavioral and photometry data were collected using custom-written workflows (Bonsai V3.5.2), processed using custom-written Python scripts (V3.7.7), and analyzed using Python and GraphPad Prism (V9.0). Transitions between chambers and cumulative time spent in each chamber were detected via binary region analysis in Bonsai and analyzed in Python. Heroin preference was calculated as the difference in time spent between chambers (i.e., Heroin preference = [time in heroin-paired] – [time in saline-paired]), and final preference was calculated as the change in preference across test sessions (i.e., CPP score = [Heroin preference, post-conditioning] – [Heroin preference, pre-conditioning]). Change in preference was analyzed using two-way repeated-measures (RM) ANOVA (dose × test) or two-tailed paired *t*-tests, and final preference was analyzed using one-way ANOVA. Equal numbers of female and male rats were included in each experiment, and sex differences in final preference were analyzed using two-way ANOVA (sex × dose) or two-tailed unpaired *t*-tests. Fos^+^ cell counts along the rostral-caudal axis of the NAc were averaged into a single value per rat and analyzed using one-way ANOVAs or two-tailed unpaired *t*-tests for each subregion. Photometry signals were de-interleaved and corrected for photobleaching by fitting the isosbestic signal (415 nm) with a first-order exponential decay curve that was scaled and subtracted from experimental traces (Proulx et al., 2018; Martianova et al., 2019).

Linearized photometry traces were then converted to z-scores (F_n_ – F_mean_ / F_SD_) using a rolling window of 10 s centered around F_n_, and events were identified where *z* > 2 (*p* < .05) with a minimum inter-event interval of 1.5 s. Photometry signals during conditioning sessions were processed (event frequency, average amplitude, variance of amplitudes) and analyzed using two-way RM ANOVA (session × drug) and Kolmogorov-Smirnov tests (cumulative distribution of amplitudes). Photometry signals during test sessions were aligned to transitions between chambers, and the window centered around each transition (−3 s to +3 s) was extracted for analysis. Transition windows were analyzed using two-way RM ANOVA (time × test), the mean signal during transitions of each type were analyzed using two-tailed paired *t*-tests, and the cumulative signal during the transition window (area under the curve [AUC]) was calculated using trapezoidal numerical integration and analyzed using two-tailed paired *t*-tests. Statistical significance for all analyses was set at *p* < .05, and all ANOVAs were followed by Sidak *post-hoc* tests (behavioral and Fos data) or Benjamini and Hochberg FDR tests with *q* < .05 (photometry data). Data are shown throughout as individual subjects and/or mean ± SEM. Subjects with lack of viral expression (*n* = 10), incorrect fiber placement (*n* = 4), or no change in preference for the heroin-paired chamber (*n* = 5) were excluded from fiber photometry experiments.

## RESULTS

### Expression of heroin CPP is accompanied by robust activation of the NAc

Female and male rats underwent a heroin CPP procedure that included an initial preference test, four days of conditioning, and a final preference test (**Figure 1*A***). To identify a dose of heroin that would reliably produce a CPP, rats were conditioned with 0, 1, or 3 mg/kg heroin. A two-way RM ANOVA revealed a significant dose × test interaction on preference for the heroin-paired chamber (*F*_(2,21)_ = 5.40, *p* = .013), with rats conditioned with 3 mg/kg heroin significantly increasing time spent in the heroin-paired chamber on the post-test compared to the pre-test (*p* = .023; **Figure 1*B***). Moreover, a one-way ANOVA revealed a significant main effect of dose on final preference (*F*_(2,21)_ = 5.28, *p* = .014), with rats conditioned with 3 mg/kg heroin (*p* = .0091) but not 1 mg/kg heroin (*p* = .078) developing a stronger CPP for the heroin-paired chamber than rats conditioned with 0 mg/kg heroin (**Figure 1*C***). To assess the role of the NAc in expression of a heroin CPP, rats were euthanized 30 min after the final preference test and brains were processed for Fos immunohistochemistry. An unpaired *t*-test revealed a significant effect of dose on Fos activation in the NAc core (*t*_(13)_ = 3.54, *p* = .0036), with significantly greater activation in rats conditioned with 3 mg/kg heroin (**Figure 1*D*– 1*E***). Similarly, an unpaired *t*-test revealed a significant effect of dose on Fos activation in the NAc shell (*t*_(13)_ = 2.94, *p* = .012), with significantly greater activation in rats conditioned with 3 mg/kg heroin (**Figure 1*F* – 1*G***). Importantly, two-way ANOVAs revealed no main effect of sex on final preference for the heroin-paired chamber (*F*_(1,18)_ = 2.46, *p* = .13) or Fos activation in the NAc core (*F*_(1,11)_ = 0.0032, *p* = .96) or NAc shell (*F*_(1,11)_ = 2.17, *p* = .17).

**Figure 1.**
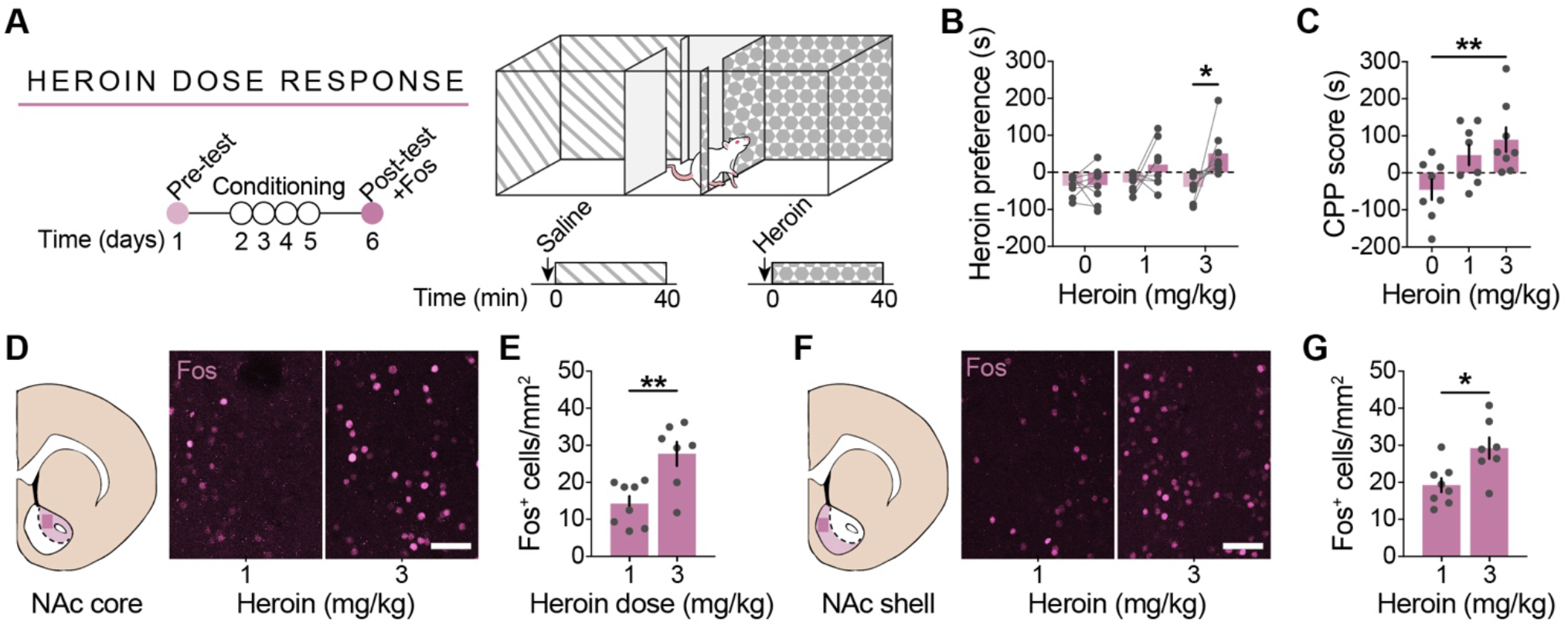
Expression of heroin CPP is accompanied with NAc activation. ***A***, Timeline for heroin CPP procedure. Rats underwent two preference tests (15 min each) separated by eight conditioning sessions (2x/day, 40 min each), and activation of the NAc during the post-test was assessed via Fos immunohistochemistry. ***B-C***, Rats conditioned with 3 mg/kg heroin spent more time in the heroin-paired chamber during the post-test and developed a significant preference for the heroin-paired chamber. ***D-E***, Representative images and quantification of Fos in the NAc core. ***F-G***, Representative images and quantification of Fos in the NAc shell. *n* = 8/group; scale bar = 50 *μ*m; \**p* < .05, \*\**p* < .01

### *In vivo* imaging of DA activity and MSN calcium activity in the NAc

To examine the role of NAc activity in the development and expression of a heroin CPP with temporal precision, CPP was coupled with fiber photometry for *in vivo* monitoring of DA signals, dMSN Ca^2+^ signals, and iMSN Ca^2+^ signals. Photometry signal quality was verified prior to starting CPP via brief home cage recordings from pairs of rats (**Figure 2*A* – 2*B***). Recordings were performed using either single-color photometry for DA imaging or dual-color photometry for simultaneous dMSN/iMSN Ca^2+^ imaging, with the isosbestic signal (415 nm) interleaved to correct for motion artifacts (**Figure 2*C***). DA imaging was performed with the DA sensor dLight1.3b (Patriarchi et al., 2018), a modified D1-type DA receptor with a cpGFP insert that expresses proximally to TH^+^ DA terminals in the NAc (**Figure 2*D* – 2*F***). Dual dMSN/iMSN Ca^2+^ imaging was performed with the green-shifted and red-shifted Ca^2+^ indicators GCaMP6s and RCaMP1b, respectively, which were selectively targeted to dMSNs and iMSNs by infusing different retrogradely transported recombinases into the VTA (AAVrg-flpo to target dMSNs) and VP (CAV2-Cre to target iMSNs; **Figure 2*G***). GCaMP6s and RCaMP1b expression was detected throughout the NAc, with limited co-expression anywhere along the rostral/caudal axis (**Figure 2*H* – 2*I***). While the majority of rats co-expressed both indicators in the NAc (*n* = 8), a minority exclusively expressed GCaMP6s (*n* = 4) or RCaMP1b (*n* = 2); thus, for all MSN Ca^2+^ imaging experiments, GCaMP6s and RCaMP1b signals were independently analyzed. An overview of the analysis pipeline used for photometry signals is shown in **Figure 2*J***. Raw signals were de-interleaved to separate experimental signals from the isosbestic signal, which was then used to linearize experimental signals and remove motion artifacts. Linearized signals were converted to normalized z-scores using a rolling mean algorithm, and significant upward deviations from the mean (“events”) were detected in normalized signals and extracted for additional analysis.

**Figure 2.**
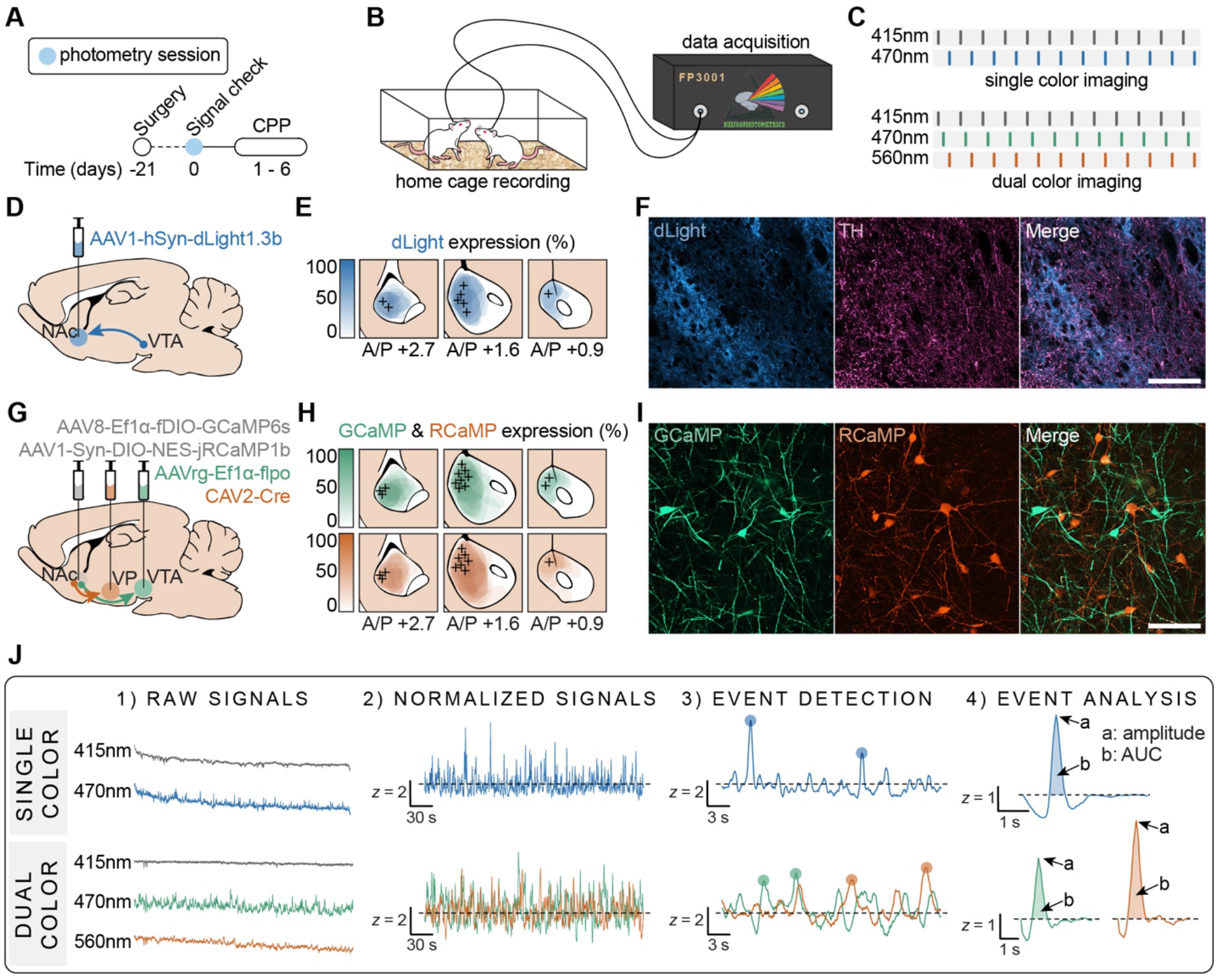
Experimental design for fiber photometry recordings in the NAc. ***A***, Timeline for photometry recordings prior to CPP. Rats underwent brief (~5 min) recordings in their home cages to verify signal quality. ***B***, Experimental setup. Rats were connected to a fiber photometry system via a branching fiber optic patch cord for *in vivo* recordings. ***C***, LED configuration for single or dual color imaging with interleaved isosbestic wavelength. ***D-E***, Viral strategy for dopamine imaging and expression of dLight1.3b throughout the NAc. ***F***, Representative dLight expression with TH staining of DA terminals in the NAc. ***G-H***, Viral strategy for dMSN/iMSN imaging and expression of GCaMP6s and RCaMP1b throughout the NAc. ***I***, Representative GCaMP and RCaMP expression in the NAc. ***J***, Analysis pipeline for photometry data. Raw signals were de-interleaved to separate channels, then linearized and converted to z-scores. Events were detected and extracted from normalized signals, and event characteristics were examined. +: optic fiber location; TH: tyrosine hydroxylase; scale bar = 100 *μ*m

### The effects of heroin on DA, dMSN, and iMSN signaling change over the course of conditioning

After confirming the quality of photometry signals, rats underwent heroin CPP with a 3 mg/kg conditioning dose. NAc DA signals (*n* = 9), NAc dMSN Ca^2+^ signals (*n* = 12), or NAc iMSN Ca^2+^ signals (*n* = 10) were recorded at four timepoints during conditioning: the saline and heroin pairings on the first day of conditioning (session 1), and the saline and heroin pairings on the fourth day of conditioning (session 4; **Figure 3*A* – 3*B***). For these recordings, pairs of rats were injected with saline or heroin, connected to the photometry system via a branching fiber optic patch cord, and immediately placed into CPP chambers. As recordings were not started until ~1min after rats received *ip* injections, analysis of photometry data was done with a rolling window across the entirety of the conditioning session, rather than using a pre-injection baseline period as a reference. A two-way RM ANOVA revealed a significant session × drug interaction on the frequency of DA events (*F*_(1,8)_ = 16.00, *p* = .004), with fewer DA events after heroin in session 1 (saline 1 vs heroin 1, *p* = .017) but more DA events after heroin in session 4 (saline 4 vs heroin 4, *p* = .008; heroin 1 vs heroin 4, *p* = .0007; **Figure 3*C* – 3*D***). Kolmogorov-Smirnov tests on the cumulative distribution of event amplitudes in each session revealed significant heroin-induced leftward shifts in both session 1 (*D* = 0.18, *p* < .0001) and session 4 (*D* = 0.25, *p* < .0001), indicative of heroin-induced suppression of large amplitude DA events (**Figure 3*E***). Indeed, two-way RM ANOVAs revealed significant main effects of drug on both the mean amplitude of DA events (*F*_(1,8)_ = 25.14, *p* = .001) and the variance of DA event amplitudes (*F*_(1,8)_ = 10.97, *p* = .011), with reduced amplitude (session 1, *p* = .017; session 4, *p* = .0085) and variance (session 1, *p* = .0017; session 4, *p* = .008) of DA events after heroin (**Figure 3*F* – 3*G***).

**Figure 3.**
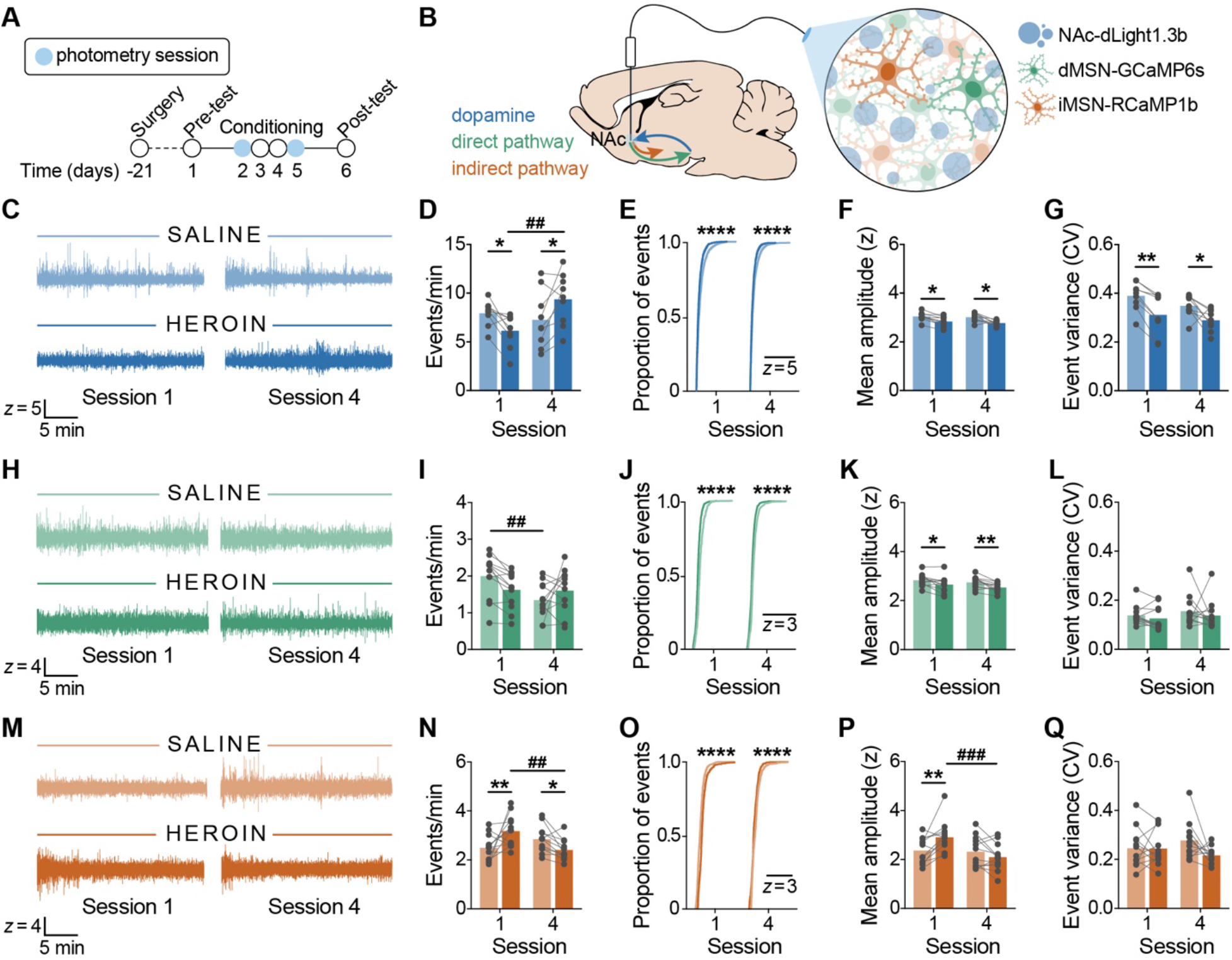
Heroin conditioning disrupts DA, dMSN, and iMSN signaling in the NAc. ***A-B***, Timeline and experimental design for photometry recordings during CPP conditioning sessions. ***C*, *H*, *M***, Representative signals collected during conditioning. ***C-D***, Heroin reduced the frequency of DA events in session 1 but increased the frequency of DA events in session 4. ***E-G***, Heroin reduced the frequency of large amplitude DA events, the average amplitude of DA events, and the variance of DA event amplitudes. ***H-I***, The frequency of dMSN events after the fourth saline pairing was significantly lower than after the first saline pairing. ***J-L***, Heroin reduced the proportion of large dMSN events and the average amplitude of dMSN events but had no effect on the variance of dMSN events. ***M-N***, Heroin increased the frequency of iMSN events in session 1 but reduced the frequency of iMSN events in session 4. ***O-P***, Heroin increased the proportion of large amplitude iMSN events and the average amplitude of iMSN events in session 1 but decreased the proportion of large amplitude iMSN events and the average amplitude of iMSN events in session 4. ***Q***, The variance of iMSN events was unchanged during conditioning. *n* = 9-12 rats; \**p* < .05, \*\**p* < .01, \*\*\**p* < .001, \*\*\*\**p* < .0001 (saline vs heroin); ^##^*p* < .01, ^###^*p* < .001 (session 1 vs session 4)

A two-way RM ANOVA on the frequency of dMSN Ca^2+^ events revealed a significant session × drug interaction (*F*_(1,11)_ = 12.96, *p* = .0042) with a selective reduction in dMSN signaling events during the fourth saline session (saline 1 vs saline 4, *p* = .0016), suggesting heroin-induced disruption in the basal level of dMSN Ca^2+^ signaling (**Figure 3*H* – 3*I***). Kolmogorov-Smirnov tests on the cumulative distribution of dMSN event amplitudes revealed significant heroin-induced leftward shifts in both session 1 (*D* = 0.53, *p* < .0001) and session 4 (*D* = 0.46, *p* < .0001), indicative of heroin-induced suppression of large amplitude dMSN signaling events (**Figure 3*J***). Accordingly, a two-way RM ANOVA revealed a significant main effect of drug on the mean amplitude of dMSN events (*F*_(1,11)_ = 17.49, *p* = .0015) but not the variance of dMSN event amplitudes (*F*_(1,11)_ = 1.03, *p* = .33), with significant heroin-induced reductions in amplitude in both session 1 (*p* = .011) and session 4 (*p* = .0043; **Figure 3*K* – 3*L***).

A two-way RM ANOVA on the frequency of iMSN Ca^2+^ events revealed a significant session × drug interaction (*F*_(1,9)_ = 19.11, *p* = .0018), with an increase in iMSN events after heroin in session 1 (saline 1 vs heroin 1, *p* = .0073) but a decrease in iMSN events after heroin in session 4 (saline 4 vs heroin 4, *p* = .023; heroin 1 vs heroin 4, *p* = .0009; **Figure 3*M* – 3*N***). Moreover, Kolmogorov-Smirnov tests on the cumulative distribution of iMSN event amplitudes revealed a heroin-induced rightward shift in session 1 (*D* = 0.49, *p* < .0001), but a leftward shift in session 4 (*D* = 0.42, *p* < .0001; **Figure 3*O***). A two-way RM ANOVA on the mean amplitude of iMSN events revealed both a significant main effect of session (*F*_(1,9)_ = 8.37, *p* = .018) and a significant session × drug interaction (*F*_(1,9)_ = 14.76, *p* = .004), with greater iMSN amplitudes after heroin in session 1 (saline 1 vs heroin 1, *p* = .0034) but weaker iMSN amplitudes after heroin in session 4 (heroin 1 vs heroin 4, *p* < .0001; **Figure 3*P***). However, two-way RM ANOVAs on the variance of iMSN event amplitudes found no significant main effect of drug (*F*_(1,9)_ = 1.76, *p* = .22) and no significant session × drug interaction (*F*_(1,9)_ = 2.07, *p* = .18; **Figure 3*Q***).

### NAc DA signaling increases when entering and decreases when exiting a heroin-paired context

Photometry signals were also recorded during both the initial (“pre-test”) and final (“post-test”) preference tests to assess the role of NAc DA signaling in the expression of a heroin CPP (**Figure 4*A* – 4*B***). A paired *t*-test revealed a significant increase in preference for the heroin-paired chamber during the post-test compared to the pre-test (*t*_(8)_ = 5.35, *p* = .0007; **Figure 4*C***), and an unpaired *t*-test found no sex difference in final preference for the heroin-paired chamber (*t*_(7)_ = 0.51, *p* = .62; data not shown). Transitions between the center chamber and the saline- and heroin-paired chambers were timestamped and aligned to photometry signals, and the window centered around each transition (−3 s to +3 s) was isolated for analysis. A two-way RM ANOVA revealed a significant time × test interaction during entries to the saline-paired chamber (*F*_(119,952)_ = 1.30, *p* = .023), with significantly weaker DA signaling in the 0.5 s centered around entry in the post-test (*p* < .008; **Figure 4*D***). Moreover, paired *t*-tests found significantly weaker DA signaling at the moment of entry (*t*_(8)_ = 4.62, *p* = .0017) as well as across the entire entry window (*t*_(8)_ = 3.94, *p* = .0043) in the post-test (**Figure 4*E* – 4*F***). However, a two-way RM ANOVA revealed no significant time × test interaction during exits from the saline-paired chamber (*F*_(119,952)_ = 1.12, *p* = .19), and paired *t*-tests found no significant difference in DA signaling at the moment of exit (*t*_(8)_ = 0.63, *p* = .55) or across the exit window (*t*_(8)_ = 0.31, *p* = .76) in the post-test (**Figure 4*G* – 4*I***). Conversely, a two-way RM ANOVA revealed a significant time × test interaction during entries to the heroin-paired chamber (*F*_(119,952)_ = 13.01, *p* < .0001), with significantly greater DA signaling in the 1s centered around entry in the post-test (*p* < .008; **Figure 4*J***). Additionally, paired *t*-tests found significantly larger DA signals at the moment of entry to the heroin-paired chamber (*t*_(8)_ = 6.70, *p* = .0002) as well as across the entire entry window (*t*_(8)_ = 6.47, *p* = .0002) in the post-test (**Figure 4*K* – 4*L***). Finally, a two-way RM ANOVA revealed a significant time × test interaction during exits from the heroin-paired chamber (*F*_(119,952)_ = 3.04, *p* < .0001), with significantly stronger DA signaling ~2 s preceding exit (*p* < .004) but significantly weaker DA signaling in the 1 s centered around exit (*p* < .003) in the post-test (**Figure 4*M***). Paired *t*-tests also found significantly weaker DA signals at the moment of exit from the heroin-paired chamber (*t*_(8)_ = 7.19*, p* < .0001) but no difference in the DA signal across the entire exit window (*t*_(8)_ = 0.0064, *p* > 0.99) in the post-test (**Figure 4*N* – 4*O***).

**Figure 4.**
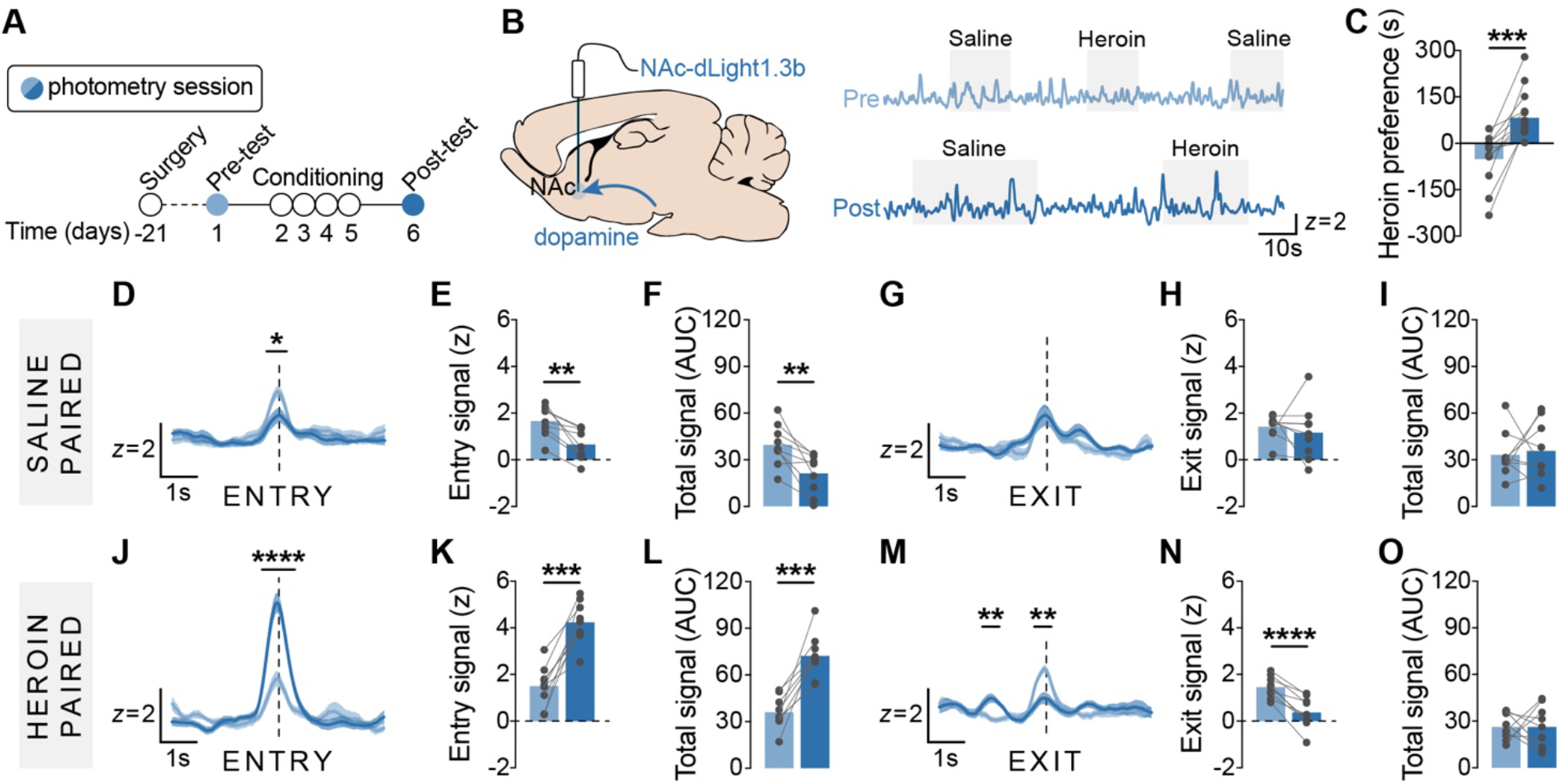
Increased DA signaling precedes entry to a heroin-paired context. ***A-B***, Timeline and strategy for NAc DA recordings during CPP test sessions, and representative DA traces collected during the pre- and post-test. ***C***, Rats significantly increased preference for the heroin-paired chamber after conditioning. ***D-O***, NAc DA signals aligned to transitions in the CPP chamber. Following conditioning, ***D-F***, DA signaling was significantly attenuated when entering the saline-paired chamber and ***G-I***, unchanged when exiting the saline-paired chamber. Following conditioning, ***J-L***, DA signaling was significantly enhanced when entering the heroin-paired chamber and ***M-O***, significantly attenuated when exiting the heroin-paired chamber. *n* = 9 rats; \**p* < .05, \*\**p* < .01, \*\*\**p* < .001, \*\*\*\**p* < .0001

### NAc dMSN signaling increases when entering and decreases when exiting a heroin-paired context

Photometry signals were also recorded during the preference tests to assess the role of NAc dMSN signaling in the expression of a heroin CPP (**Figure 5*A* – 5*B***). A paired *t*-test revealed a significant increase in preference for the heroin-paired chamber during the post-test compared to the pre-test (*t*_(11)_ = 3.17, *p* = .009; **Figure 5*C***), and an unpaired *t*-test found no sex difference in final preference for the heroin-paired chamber (*t*_(10)_ = 0.16, *p* = .88; data not shown). A two-way RM ANOVA revealed a significant time × test interaction during entries to the saline-paired chamber (*F*_(79,869)_ = 5.26, *p* < .0001), with significantly weaker dMSN signaling from 1 s preceding to 1.5 s following entry (*p* < .008) in the post-test (**Figure 5*D***). Moreover, paired *t*-tests found significantly weaker dMSN signaling at the moment of entry (*t*_(11)_ = 5.83, *p* = .0001) as well as across the entire entry window (*t*_(11)_ = 3.26, *p* = .0076) in the post-test (**Figure 5*E* – 5*F***). However, a two-way RM ANOVA revealed no significant time × test interaction during exits from the saline-paired chamber (*F*_(79,869)_ = 1.27, *p* = .063), and paired *t*-tests found no significant difference in dMSN signaling at the moment of exit (*t*_(11)_ = 0.20, *p* = .84) or across the exit window (*t*_(11)_ = 0.069, *p* = .95) in the post-test (**Figure 5*G* – 5*I***). Conversely, a two-way RM ANOVA revealed a significant time × test interaction during entries to the heroin-paired chamber (*F*_(79,869)_ = 18.71, *p* < .0001), with significantly greater dMSN signaling in the 2 s centered around entry in the post-test (*p* < .002; **Figure 5*J***). Additionally, paired *t*-tests found significantly larger dMSN signals at the moment of entry to the heroin-paired chamber (*t*_(11)_ = 9.34, *p* < .0001) as well as across the entire entry window (*t*_(11)_ = 3.87, *p* = .0026) in the post-test (**Figure 5*K* – 5*L***). Finally, a two-way RM ANOVA revealed a significant time × test interaction during exits from the heroin-paired chamber (*F*_(79,869)_ = 5.78, *p* < .0001), with significantly weaker dMSN signaling in the 1 s centered around exit (*p* < .002) in the post-test (**Figure 5*M***). Paired *t*-tests also found significantly weaker dMSN signals at the moment of exit from the heroin-paired chamber (*t*_(11)_ = 6.29*, p* < .0001) but no difference in the dMSN signal across the entire exit window (*t*_(11)_ = 2.00, *p* = 0.071) in the post-test (**Figure 5*N* – 5*O***).

**Figure 5.**
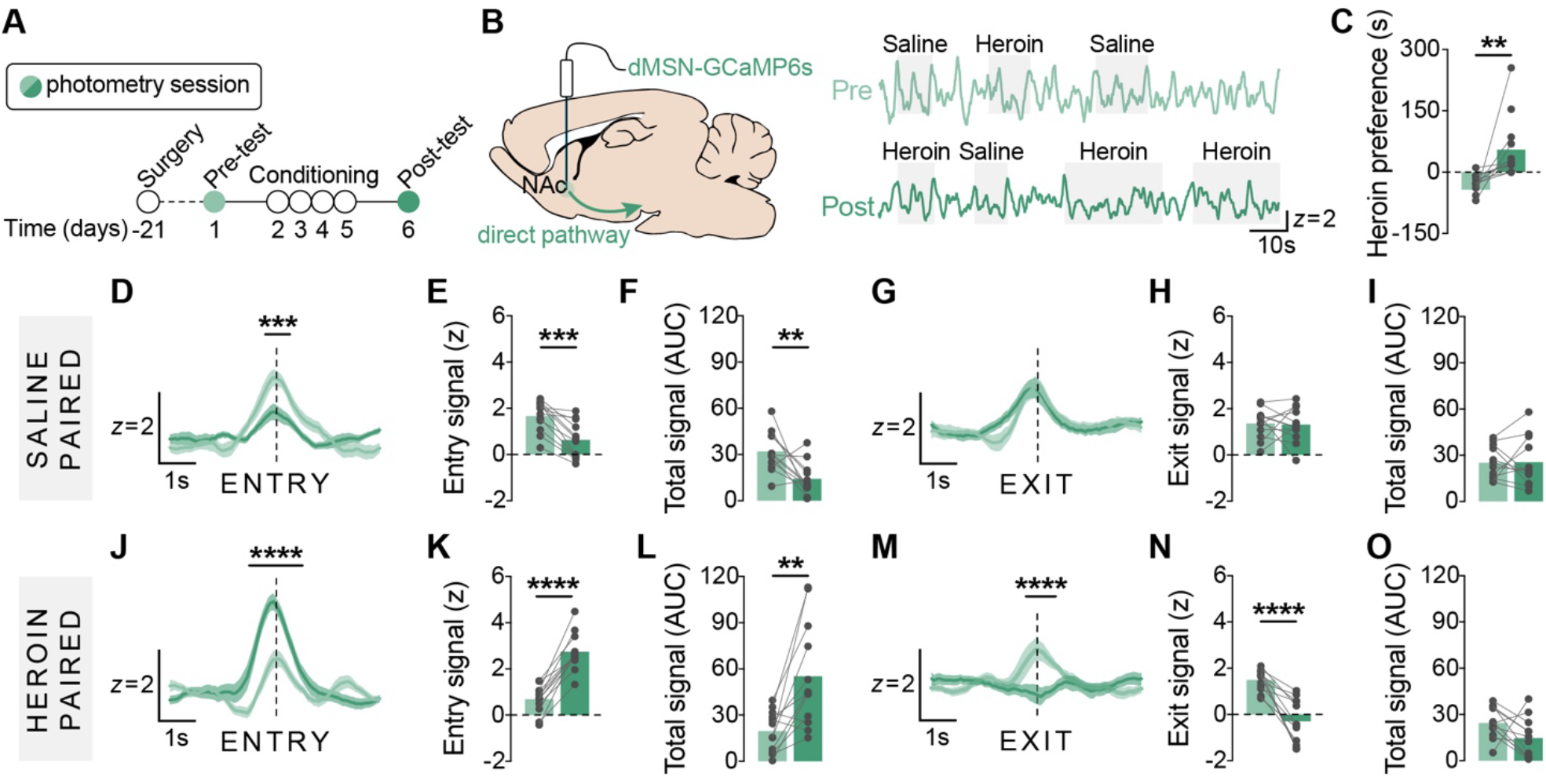
Increased dMSN signaling precedes entry to a heroin-paired context. ***A-B***, Timeline and strategy for NAc dMSN recordings during CPP test sessions, and representative dMSN traces collected during the pre- and post-test. ***C***, Rats significantly increased preference for the heroin-paired chamber after conditioning. ***D-O***, NAc dMSN signals aligned to transitions in the CPP chamber. Following conditioning, ***D-F***, dMSN signaling was significantly attenuated when entering the saline-paired chamber and ***G-I***, unchanged when exiting the saline-paired chamber. Following conditioning, ***J-L***, dMSN signaling was significantly enhanced when entering the heroin-paired chamber and ***M-O***, significantly attenuated when exiting the heroin-paired chamber. *n* = 12 rats; \*\**p* < .01, \*\*\**p* < .001, \*\*\*\**p* < .0001

### NAc iMSN signaling decreases when entering and increases when exiting a heroin-paired context

Photometry signals were also recorded during the preference tests to assess the role of NAc iMSN signaling in the expression of a heroin CPP (**Figure 6*A* – 6*B***). A paired *t*-test revealed a significant increase in preference for the heroin-paired chamber during the post-test compared to the pre-test (*t*_(10)_ = 3.28, *p* = .0084; **Figure 6*C***), and an unpaired *t*-test found no sex difference in final preference for the heroin-paired chamber (*t*_(8)_ = 0.42, *p* = .68; data not shown). A two-way RM ANOVA revealed a significant time × test interaction during entries to the saline-paired chamber (*F*_(79,711)_ = 2.88, *p* < .0001), with significantly stronger iMSN signaling from 0.5 s preceding to 1.5 s following entry (*p* < .006) in the post-test (**Figure 6*D***). Moreover, paired *t*-tests found significantly greater iMSN signaling at the moment of entry (*t*_(9)_ = 3.50, *p* = .0067) as well as across the entire entry window (*t*_(9)_ = 2.32, *p* = .045) in the post-test (**Figure 6*E* – 6*F***). However, a two-way RM ANOVA revealed no significant time × test interaction during exits from the saline-paired chamber (*F*_(79,711)_ = 0.67, *p* = .99), and paired *t*-tests found no significant difference in iMSN signaling at the moment of exit (*t*_(9)_ = 1.33, *p* = .22) or across the exit window (*t*_(9)_ = 2.23, *p* = .053) in the post-test (**Figure 6*G* – 6*I***). Conversely, a two-way RM ANOVA revealed a significant time × test interaction during entries to the heroin-paired chamber (*F*_(79,711)_ = 2.40, *p* < .0001), with significantly weaker iMSN signaling in the 1 s centered around entry in the post-test (*p* < .003; **Figure 6*J***). Additionally, paired *t*-tests found significantly smaller iMSN signals at the moment of entry to the heroin-paired chamber (*t*_(9)_ = 2.55, *p* = .031) as well as across the entire entry window (*t*_(9)_ = 3.64, *p* = .0054) in the post-test (**Figure 6*K* – 6*L***). Finally, a two-way RM ANOVA revealed a significant time × test interaction during exits from the heroin-paired chamber (*F*_(79,711)_ = 1.63, *p* = .0008), with significantly stronger iMSN signaling in the 1 s centered around exit (*p* < .006) in the post-test (**Figure 6*M***). Paired *t*-tests also found significantly stronger iMSN signals at the moment of exit from the heroin-paired chamber (*t*_(9)_ = 3.57*, p* = .006) as well as across the entire exit window (*t*_(9)_ = 3.53, *p* = 0.0064) in the post-test (**Figure 6*N* – 6*O***).

**Figure 6.**
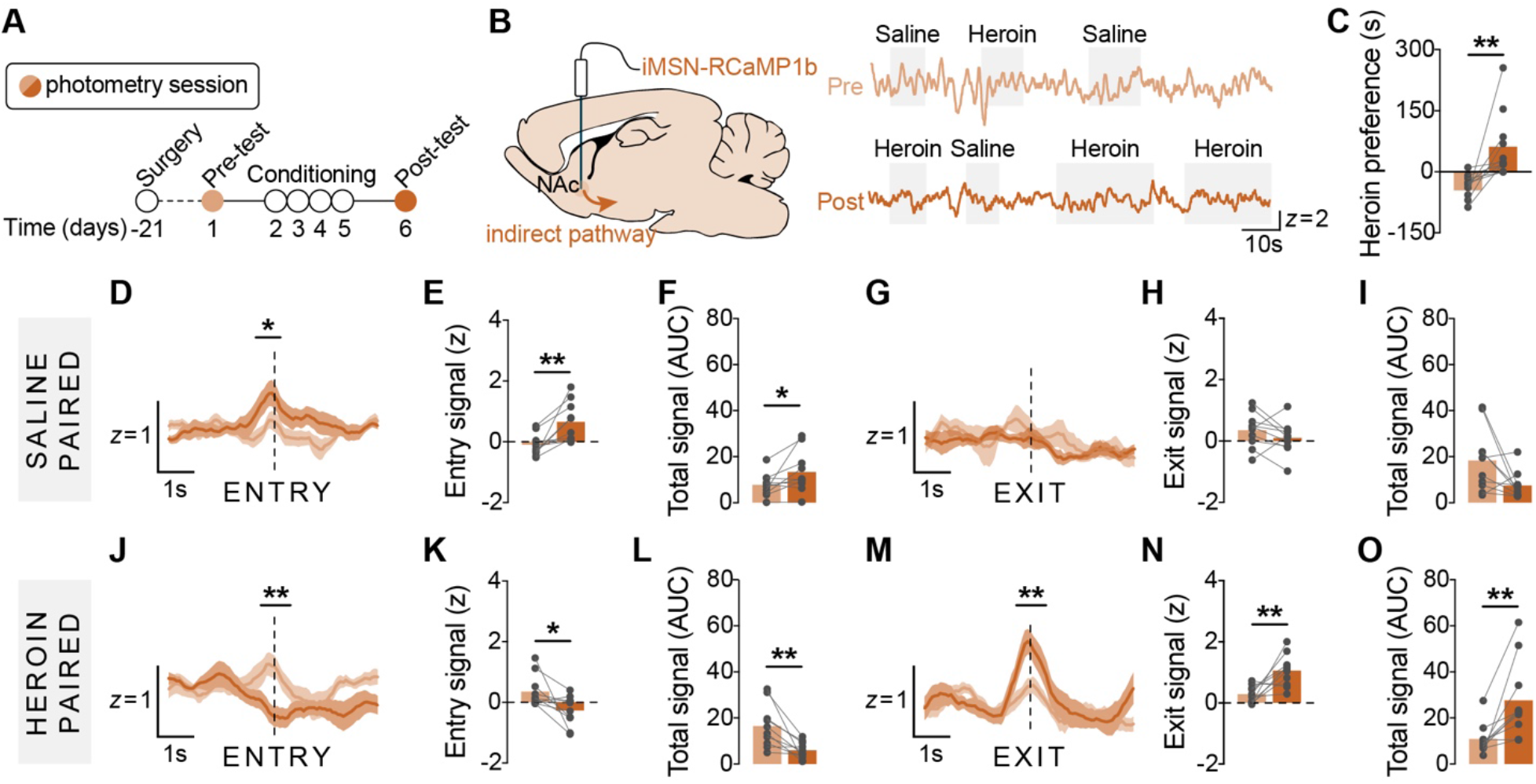
Decreased iMSN signaling precedes entry to a heroin-paired context. ***A-B***, Timeline and strategy for NAc iMSN recordings during CPP test sessions, and representative iMSN traces collected during the pre- and post-test. ***C***, Rats significantly increased preference for the heroin-paired chamber after conditioning. ***D-O***, NAc iMSN signals aligned to transitions in the CPP chamber. Following conditioning, ***D-F***, iMSN signaling was significantly enhanced when entering the saline-paired chamber and ***G-I***, unchanged when exiting the saline-paired chamber. Following conditioning, ***J-L***, iMSN signaling was significantly attenuated when entering the heroin-paired chamber and ***M-O***, significantly enhanced when exiting the heroin-paired chamber. *n* = 10 rats; \**p* < .05, \*\**p* < .01

### Buprenorphine pretreatment blocks the development of heroin CPP

Buprenorphine is used for opioid replacement therapy due to its ability to weakly activate mu opioid receptor signaling and prevent the binding of other opioids (e.g., heroin), and was recently shown to occlude heroin-evoked DA release into the NAc (Isaacs et al., 2020). To determine whether buprenorphine could similarly prevent the acquisition of heroin CPP, rats underwent a modified CPP procedure with 0 or 0.2 mg/kg buprenorphine given 10 min prior to each conditioning session (**Figure 7*A***). A two-way RM ANOVA revealed a significant dose × test interaction on heroin preference (*F*_(1,14)_ = 8.52, *p* = .012), with a significant increase in time spent in the heroin-paired chamber for rats pretreated with 0 mg/kg buprenorphine (*p* = .0086) but not 0.2 mg/kg buprenorphine (*p* = .73; **Figure 7*B***). Moreover, an unpaired *t*-test revealed significantly lower final preference for the heroin-paired chamber in rats pretreated with 0.2 mg/kg buprenorphine (*t*_(14)_ = 2.83, *p* = .014; **Figure 7*C***). To assess the impact of buprenorphine pretreatment during conditioning on NAc activation during the final preference test, rats were euthanized 30 min after the final preference test and brains were processed for Fos immunohistochemistry. Unpaired *t*-tests revealed significantly lower levels of Fos activation in both the NAc core (*t*_(14)_ = 4.78, *p* = .0003) and NAc shell (*t*_(14)_ = 6.20, *p* < .0001) for rats pretreated with 0.2 mg/kg buprenorphine during conditioning (**Figure 7*D*– 7*G***). Importantly, two-way ANOVAs revealed no main effect of sex on final preference for the heroin-paired chamber (*F*_(1,12)_ = 1.40, *p* = .26) or Fos activation in the NAc core (*F*_(1,12)_ = 1.09, *p* = .32) or NAc shell (*F*_(1,12)_ = 0.26, *p* = .62).

**Figure 7.**
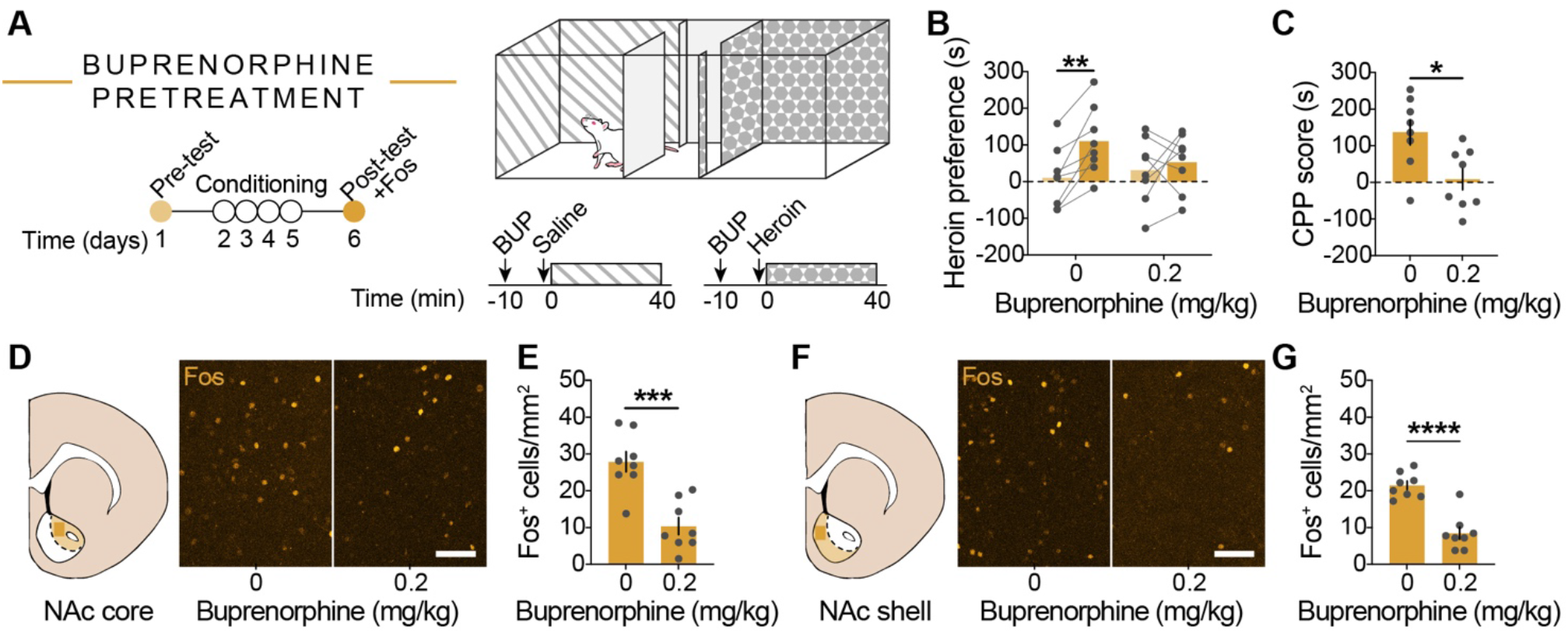
Acquisition of heroin CPP is blocked by buprenorphine. ***A***, Timeline for buprenorphine experiment. Testing was performed as described in **Figure 1**, but rats received buprenorphine pretreatment 10 min prior to each conditioning session. ***B-C***, Rats pretreated with 0.2 mg/kg buprenorphine spent significantly less time in the heroin-paired chamber during the post-test than those pretreated with 0 mg/kg buprenorphine and did not develop a preference for the heroin-paired chamber. ***D-G***, Representative images and quantification of Fos in the NAc. Rats pretreated with 0.2 mg/kg buprenorphine had significantly less Fos in both subregions of the NAc than those pretreated with 0 mg/kg buprenorphine. *n* = 8/group; scale bar = 50 *μ*m; \**p* < .05, \*\**p* < .01, \*\*\**p* < .001, \*\*\*\**p* < .0001

## DISCUSSION

Here, we sought to examine the role of NAc signaling in the acquisition and expression of heroin CPP by coupling fiber photometry recordings with a heroin CPP procedure that induces Fos activation in the NAc during expression of heroin CPP. We identified differences in the neural response to heroin over the course of conditioning, with suppression of NAc activity during early conditioning but enhancement of NAc activity during late conditioning. Next, we demonstrated that entering a heroin-paired context is accompanied by increased dopamine and dMSN Ca^2+^ signaling along with decreased iMSN Ca^2+^ signaling, while exiting a heroin-paired context is accompanied by increased iMSN signaling along with decreased dopamine and dMSN signaling. Finally, we show that buprenorphine pretreatment during conditioning is sufficient to prevent the expression of heroin CPP and concomitant activation of the NAc. Together, these data reveal a central role for the NAc in the reinforcing effects of heroin, in accordance with the hypothesis that an imbalance in signaling between the accumbal direct and indirect pathways drives addictive behaviors.

### Dopaminergic modulation in the NAc contributes to heroin conditioning

While the contribution of DA to the rewarding effects of most potentially addictive drugs (e.g., psychostimulants, nicotine) is well-established (Crummy et al., 2020), the role of DA in opioid reward is more complicated (Badiani et al., 2011). Opioids increase phasic DA release, and both D1 receptor blockade and D2 receptor deletion blocks morphine reward (Johnson and North, 1992; Shippenberg et al., 1993; Maldonado et al., 1997; Sellings and Clarke, 2003). However, bilateral 6-OHDA lesions of the NAc can block or have no effect on morphine reward (Shippenberg et al., 1993; Sellings and Clarke, 2003), and chronic blockade of both D1 and D2 receptors potentiates the rewarding effects of low doses of heroin (Stinus et al., 1989). During conditioning, we identified noteworthy effects of heroin on DA signaling: a significant reduction in large amplitude DA events during both early and late conditioning sessions, a significant reduction in the frequency of DA events during early conditioning, and a significant enhancement in the frequency of DA events during late conditioning. This shift from suppression to elevation across conditioning suggests sensitization of the dopaminergic response to heroin and supports a role for DA in the acquisition of heroin CPP. After conditioning, we detected a significant increase in DA signaling preceding entries to the heroin-paired chamber, as well as significant decreases in DA signaling preceding exits from the heroin-paired chamber and entries to the saline-paired chamber, indicating emergent selectivity for DA signaling in response to a heroin-paired context. Notably, a recent study using DA imaging in the NAc medial shell following heroin administration reported a significant rise in tonic DA signaling over a period of minutes, relative to a pre-injection baseline (Corre et al., 2018). We did not observe a similar effect on DA signaling during conditioning, although we believe this is largely due to lack of a pre-injection baseline in the study design. Future work will expand upon this data to better understand how heroin conditioning is altering tonic signaling in the NAc, in addition to phasic changes.

### An imbalance in accumbal activity develops during heroin conditioning

The ability of DA to alter overall accumbal activity is dependent not only on dopaminergic modulation of individual MSNs but also on the network implications of such modulation. Although dMSNs and iMSNs can be differentiated according to their downstream targets (VTA and VP, respectively), both subtypes of MSNs are heavily interconnected in a local microcircuit of lateral inhibition within the NAc (Burke et al., 2017). Indeed, although dMSNs are more likely to collateralize with other dMSNs than iMSNs (Taverna et al., 2008), D2 receptor-mediated disinhibition of iMSN lateral inhibition onto dMSNs is necessary for the locomotor sensitizing effects of cocaine (Dobbs et al., 2016). An imbalance between dMSNs and iMSNs can thereby shape the ability of DA to modulate accumbal activity, with the overall balance in signaling guiding vulnerability to the reinforcing effects of drugs.

We observed a shift in the response of NAc iMSNs to heroin over the course of conditioning, with a prominence of iMSN signaling during early conditioning (enhanced iMSN frequency and amplitudes) and a suppression of iMSN signaling during late conditioning (reduced iMSN frequency and amplitudes). This inhibition of iMSN signaling – mediated in part by an enhanced dopaminergic response to heroin during late conditioning – would release nearby dMSNs from tonic lateral inhibition, allowing direct pathway signaling to dominate. Interestingly, however, we did not observe a sensitized dMSN response to heroin over the course of conditioning; rather, we observed a transient reduction in dMSN signaling during the fourth saline session, indicating a weakening of basal dMSN tone over the course of heroin conditioning. In addition to heroin-induced disruptions in NAc activity over the course of conditioning, we identified context-dependent changes in MSN signals post-conditioning. dMSN signals were significantly stronger when entering the heroin-paired context, but significantly weaker when exiting the heroin-paired or entering the saline-paired contexts. Conversely, iMSN signals were significantly weakened when entering the heroin-paired context, but significantly stronger when exiting the heroin-paired or entering the saline-paired contexts. These data indicate selective tuning of dMSNs to the heroin-paired context, and are in consistent with previous findings of elevated dMSN signaling during re-exposure to a cocaine-paired context (Calipari et al., 2016).

### Disrupting opioid signaling prevents development of heroin CPP

In our final experiment, we show that pretreatment with buprenorphine during conditioning blocked the development of a heroin CPP and associated Fos activity in the NAc. Although buprenorphine weakly activates mu opioid receptors to mildly elevate DA release, it also prevents heroin from binding to the same receptors and driving larger phasic DA release (Isaacs et al., 2020). The buprenorphine dose used in the present study was selected based on a previous report that described a bell-shaped curve for buprenorphine-stimulated DA release into the NAc: significant increases in DA release with 0.01 – 0.04 mg/kg buprenorphine, but no change in DA release with 0.18 – 0.7 mg/kg buprenorphine (Isaacs et al., 2020). Moreover, twice daily injections of 0.1 – 0.4 mg/kg buprenorphine were sufficient to significantly attenuate cocaine self-administration in rats, with no differences observed between 0.1 and 0.4 mg/kg (Carroll and Lac, 1992). Thus, our dose of buprenorphine administered prior to each conditioning session likely produced comparably low levels of DA release in both chambers, preventing the acquisition of a strong conditioned response to heroin.

### Technical considerations

Although dMSNs and iMSNs have canonically been assumed to project exclusively to the VTA and VP, respectively, growing evidence has demonstrated that dMSNs project equally strong to the VTA and the VP (Kupchik et al., 2015; Creed et al., 2016). Importantly, the viral strategy used here has been shown to target largely non-overlapping populations of neurons in the NAc with bidirectional control over cue-induced heroin-seeking (O’Neal et al., 2020). In the present study we not only did not observe co-expression of GCaMP6s and RCaMP1b within the NAc, but we also observed oppositional dMSN and iMSN signals during transitions in the post-test, similar to what has previously been reported during cocaine CPP (Calipari et al., 2016). Furthermore, although dual-color imaging runs the risk of signal crosstalk and contamination (Meng et al., 2018), our dMSN-GCaMP6s and iMSN-RCaMP1b signals were highly uncorrelated throughout testing (pre-conditioning *r^2^*: mean = 0.02, range = 3.4e^−7^ to 0.08; post-conditioning *r^2^*: mean = 0.05, range = 4.0e^−3^ to 0.17). Additionally, there is concern that the viral strategy used to target dMSNs may have led to undesired expression of GCaMP6s in DA terminals in the NAc. While retrograde AAVs have a higher tropism for axon terminals, some evidence has suggested they may also infect cell bodies at the site of infusion (Tervo et al., 2016); moreover, AAV8s have been shown to undergo some degree of retrograde transport, particularly when infused near DA terminals (Masamizu et al., 2011; Löw et al., 2013). However, using TH immunohistochemistry, we did not detect GCaMP6s expression in either DA terminals in the NAc or in DA cell bodies in the VTA (data not shown).

## Acknowledgements

This work was supported by grants from the National Institute on Drug Abuse (F31DA047012 to TJO, R01DA036582 to SMF) and the UW Addictions, Drug & Alcohol Institute (ADAI-0138 to TJO). We thank Dr. John Neumaier and Dr. Michelle Kelly for providing the CAV2-Cre virus, the UW Molecular Genetics Resource Core (P30DA048736) for producing the AAV1-dLight1.3b virus, and Dr. Elizabeth Crummy, Dr. Scott Ng-Evans, and Joshua O’Neal for programming assistance.

## Conclusions

Together, these data highlight a central role for NAc dMSN, iMSN, and DA signaling in the acquisition and expression of heroin CPP. Future work will investigate the role of NAc signaling in encoding individual vulnerability to the rewarding and motivating effects of heroin, with a particular focus on the relative strength of dMSNs and iMSNs between individuals sensitive to versus resistant to the conditioned reinforcement produced by heroin.

